# Longitudinal CCR2 PET Imaging Tracks Early Inflammatory Monocyte and Macrophage Infiltration in Experimental Arthritis

**DOI:** 10.64898/2026.11.09.765427

**Authors:** Maria I. Menendez Montes, Rajiu Venkatesan, Kristin L. Lenz, Hannah Luehmann, Gyu Seong Heo, Luke Springer, Huimin Yan, Deborah Sultan, Joan Tao, Vincent Gao, Yongjian Liu, Christine T.N. Pham, Farshid Guilak

## Abstract

**Objective:** Recruitment of C-C motif chemokine receptor 2 (CCR2^+^) inflammatory monocytes and macrophages represent an early event in rheumatoid arthritis (RA) pathogenesis that cannot be tracked noninvasively with existing imaging modalities. We evaluated longitudinal ^68^Ga-DOTA-ECL1i positron emission tomography/computed tomography (PET/CT) to track and quantify CCR2^+^ cell recruitment in experimental inflammatory arthritis.

**Methods:** Longitudinal ^68^Ga-DOTA-ECL1i PET/CT imaging was performed in K/BXN serum transfer arthritis (STA) mice. Imaging findings were validated using biodistribution, autoradiography, histopathology, immunofluorescence, microCT, cytokine profiling, clinical assessment, and pain-related behavioral testing.

**Results:** Joint ^68^Ga-DOTA-ECL1i uptake increased during the acute arthritis and declined with disease resolution. Increased tracer signal localized to inflamed joints, coinciding with CCR2^+^ monocyte and macrophage infiltration, synovial inflammation, cartilage damage, local CCL2 elevation, and mechanical allodynia. PET uptake correlated with ankle swelling, clinical disease severity, pain, and paw CCL2 levels, with peak tracer signal preceding progressive structural bone remodeling.

**Conclusion:** Longitudinal ^68^Ga-DOTA-ECL1i PET/CT enables noninvasive visualization and quantification of early CCR2^+^ monocyte and macrophage recruitment, establishing CCR2 PET imaging as a mechanism-based molecular platform for early detection, risk stratification, and therapeutic monitoring in inflammatory arthritis.

## Introduction

Rheumatoid arthritis (RA) is a systemic autoimmune disease characterized by persistent synovitis and progressive cartilage and bone destruction, leading to pain and disability [1]. Although early treatment and treat-to-target strategies improve clinical outcomes, sustained remission remains difficult to achieve [2, 3]. Clinical management currently relies primarily on indices such as the Disease Activity Score in 28 joints, Simplified Disease Activity Index (DAI), and Clinical Disease Activity Index (CDAI) [4, 5]; however, these provide indirect estimates of inflammatory activity and do not identify the pathogenic cellular processes responsible for persistent synovitis or progressive joint damage. A mechanism-based biomarker capable of detecting active inflammation at a cellular level could therefore provide information that guides personalized treatment for improved outcomes. Monocytes and macrophages are central to synovial inflammation. Increased numbers of subsynovial macrophages are an early feature of active RA, and correlate with disease activity and joint erosion [6]. Circulating inflammatory monocytes originate predominantly from the bone marrow and serve as precursors of infiltrating macrophages and osteoclasts [7]. Their recruitment into inflamed joints is regulated in part by the C-C motif chemokine ligand 2 and its receptor (CCL2–CCR2) axis. CCL2, also known as monocyte chemoattractant protein-1 (MCP-1), is produced by activated synoviocytes and establishes a chemotactic gradient for CCR2-expressing monocytes [8]. Increased CCL2 expression has been demonstrated in the blood and synovial tissues of patients with RA, supporting a role for this pathway in leukocyte recruitment and chronic synovial inflammation [9]. Following tissue infiltration, recruited CCR2^+^ monocytes can differentiate into inflammatory macrophages that produce tumor necrosis factor (TNF-α), interleukin-1β (IL-1β), interleukin-6 (IL-6), chemokines, matrix-degrading enzymes, and osteoclastogenic mediators [6, 7]. These cells thereby amplify synovitis and contribute to cartilage degradation, bone erosion, and pain.

The pathogenic relevance of CCR2 in RA is supported by extensive preclinical and clinical evidence. Contemporary reviews identify CCR2 as an important regulator of monocyte trafficking, synovial inflammation, fibroblast activation, osteoclastogenesis, and joint destruction [10]. CCR2 blockade further established that the pathway can be pharmacologically targeted in patients with RA, although receptor inhibition alone did not produce meaningful clinical improvement, potentially because of chemokine-network redundancy and compensatory inflammatory pathways [11]. This therapeutic outcome does not diminish the value of CCR2 as a biomarker. Rather, CCR2 expression may provide a means to identify and quantify a defined inflammatory cell population without assuming that inhibition of a single chemokine receptor will be sufficient to suppress the entire disease process. In this context, CCR2-targeted imaging captures monocyte and macrophage recruitment in real time leaving the broader inflammatory network intact for further analysis.

Conventional imaging provides important but incomplete information about inflammatory arthritis. Radiography can identify bone erosion and joint-space loss but is insensitive to early inflammatory changes [12, 13]. Ultrasound and magnetic resonance imaging reveal subclinical inflammation, synovitis, and bone marrow edema prior to visible radiographic damage [12-18]. However, these modalities primarily detect tissue-level abnormalities and cannot directly identify or quantify inflammatory monocytes and macrophages. Molecular imaging with PET offers an opportunity to interrogate disease biology in vivo. For example, ^18^F-fluorodeoxyglucose (FDG) PET measures indicate increased glucose metabolism in inflamed synovium and may reflect RA disease activity, but uptake is not specific for a particular immune-cell population or molecular pathway [19-21]. Translocator protein–targeted (TSPO) PET has been evaluated as an imaging marker of macrophage and stromal activation in RA pannus, but translocator protein is expressed by several activated cell populations and does not selectively identify recruited inflammatory monocytes and macrophages [22, 23]. Fibroblast activation protein inhibitor (FAPI) PET provides complementary information about activated synovial fibroblasts and has shown promise for evaluating chronic RA [24-28]. These studies establish the value of targeted molecular imaging while highlighting the need for direct tissue inflammatory monocyte and macrophage visualization.

CCR2-targeted PET addresses this unmet need by interrogating a chemokine receptor associated with inflammatory cell trafficking rather than nonspecific consequence of inflammation. In this regard, chemokine-receptor radiotracers have emerged as tools for noninvasive assessment of immune-cell localization and activity [29]. The CCR2-targeted radiotracer ^68^Ga-DOTA-ECL1i demonstrated specific detection of CCR2^+^ inflammatory cells in experimental lung transplantation and inflammatory lung disease [30, 31]. In myocardial injury, CCR2 PET visualized the recruitment of inflammatory monocytes and macrophages, enabling longitudinal evaluation of the inflammatory response [32, 33]. Clinical translation in pulmonary fibrosis demonstrated the feasibility for ^68^Cu-DOTA-ECL1i for imaging CCR2-associated inflammation in humans [34]. These findings support CCR2 PET as a translational platform for quantifying infiltrating inflammatory monocytes and macrophages, but its ability to track temporal CCR2-mediated inflammation in arthritis requires further validation.

The K/BxN serum transfer arthritis is a reproducible joint inflammation model involving innate immune cell infiltration and structural remodeling, thereby providing a relevant system for evaluating the effector phase of inflammatory arthritis [35-38]. We hypothesized that longitudinal ^68^Ga-DOTA-ECL1i PET can noninvasively visualize and quantify the early recruitment of CCR2^+^ inflammatory monocytes and macrophages into arthritic joints and that tracer uptake reflects disease progression at a cellular and molecular level. To test this hypothesis, we integrated longitudinal CCR2 PET with autoradiography, histopathology, immunofluorescence, cytokine profiling, microcomputed tomography, clinical arthritis assessment, and pain-related behavior.

## Materials and Methods

All procedures were conducted in accordance with protocols approved by the Washington University in St. Louis Institutional Animal Care and Use Committee and adhered to the ARRIVE guidelines.

### K/BxN model of inflammatory arthritis

The model and methods were adapted from previous work [39-41]. Male 6–8-week-old C57BL/6 mice obtained from The Jackson Laboratory (Bar Harbor, ME) were administered 200 µL of K/BxN serum by retro-orbital injection to induce arthritis. PET imaging was performed prior to K/BxN serum transfer arthritis (STA) induction, and 7-, 14-, and 23-days post-K/BxN STA (Figure 1). Disease activity (clinical score and ankle thickness) and pain behavior (electronic Von Frey) were assessed throughout the study (Figure 2). The clinical score was assessed on a scale of 0 to 3 (0, no swelling or erythema; 1, slight swelling or erythema; 2, moderate erythema and swelling in multiple digits or entire paw; and 3, pronounced erythema and swelling of entire paw; maximum total score of 12). The change from baseline in ankle thickness was determined by dial calipers, and an average change in the ankle thickness was determined for each mouse from the two hind paw measurements [27-29]. Mice were sacrificed at day 25 post disease induction and serum, front and hind paws were collected for analysis.

**Figure 1.**
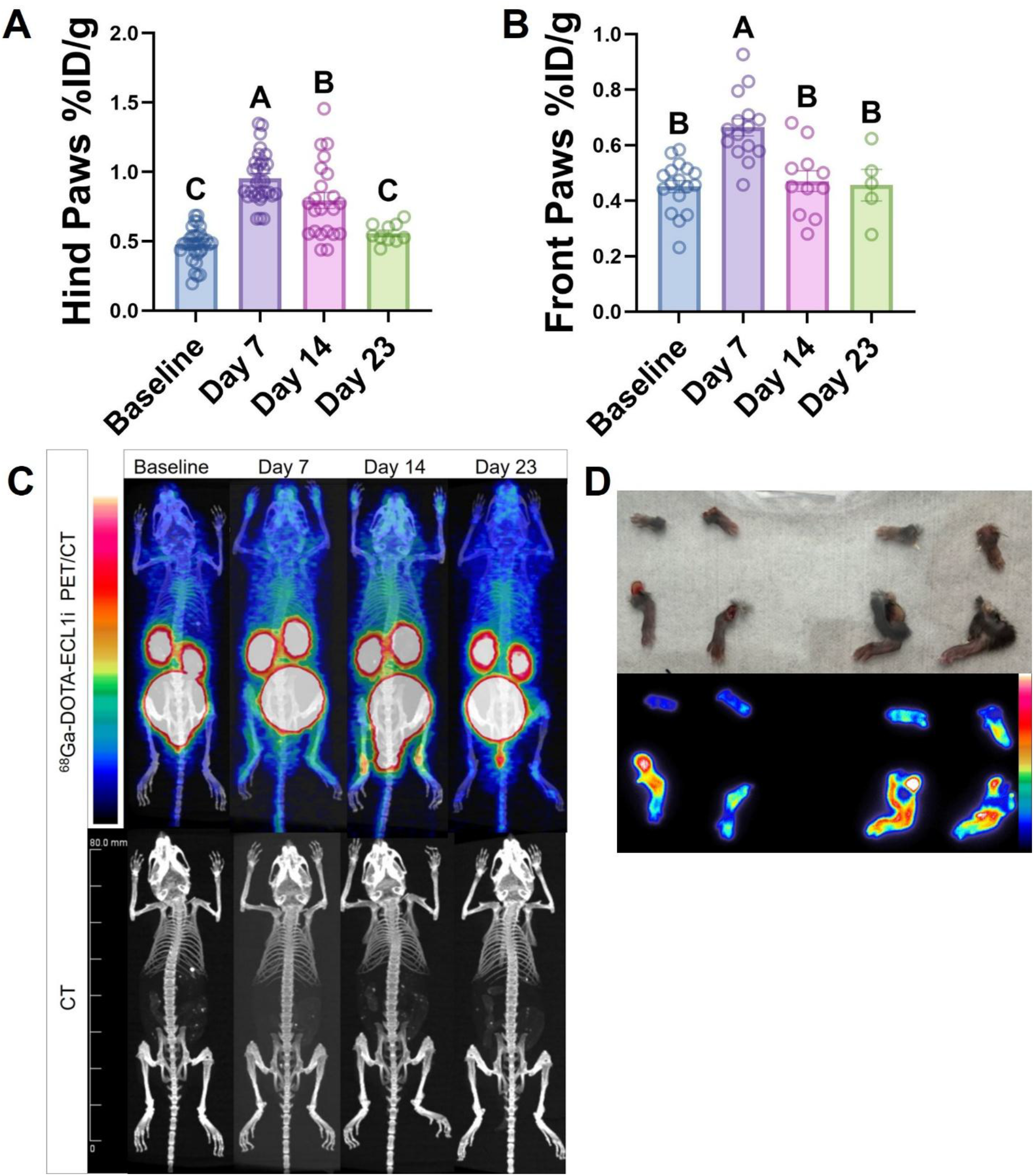
^68^Ga-DOTA-ECL1i PET imaging of CCR2 expression in the K/BxN STA model. Quantitative PET analysis of the front and hind paws (*n* = 5–17/group) demonstrated increased tracer uptake in the hind paws at days 7 and 14 and in the front paws at day 7 following K/BxN STA induction (A). Representative maximum intensity projection (MIP) fused PET/CT and CT images acquired at baseline and days 7, 14, and 23 after K/BxN STA induction are shown (B). Representative autoradiographs of the front and hind paws at day 14 are shown (D). Data are presented as mean ± SEM. Different letters indicate significant differences (*P* < 0.05) determined by one-way ANOVA with Tukey’s multiple-comparison test (A).

**Figure 2.**
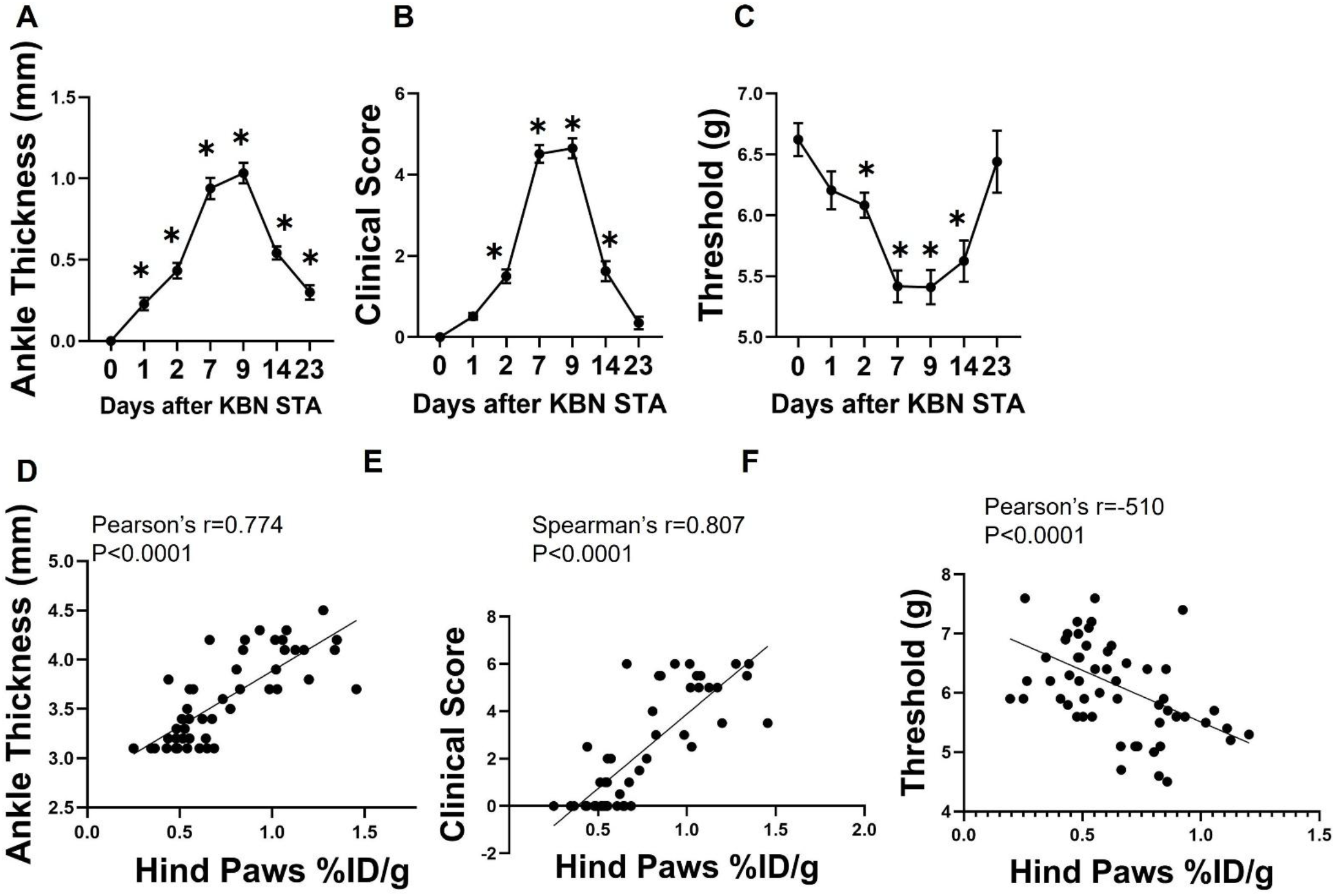
Clinical assessment of arthritis progression in the K/BxN STA model. Ankle thickness (A), combined clinical score of the front and hind paws (B), and tactile allodynia measured by the electronic von Frey assay (C) were assessed longitudinally following K/BxN STA induction (*n* = 5–18/group). Data are presented as mean ± SEM. Statistical analysis was performed using one-way ANOVA with Dunnett’s multiple-comparison test. *P* < 0.05 versus baseline.

### PET acquisition and image analysis

Mice underwent PET/CT imaging scans at baseline, and at 7, 14, and 23 days post-K/BxN STA induction. Mice received a tail intravenous injection of ≃ 3.7-6 megabecquerel (MBq) of ^68^Ga-DOTA-ECL1i. Whole-body PET/ CT scans were performed at 45–60 min (3 frames, 5 min/frame) post injection. PET images were reconstructed with the maximum a posteriori algorithm and processed using the Inveon Research Workplace (IRW) software (Siemens). All three frames were summed to acquire an average PET signal over the 15 min scan period. Three-dimensional volumes of interest (VOI) with a total volume of 6 mm^3^ were drawn on the PET images over the front and hind paws of each mouse. Uptake values for each ROI were generated by the IRW software and converted to percent injected dose per gram of tissue (%ID/g) using a tissue density of 1 g/mm^3^ and the amount of radioactivity injected into the mouse decay corrected to the scan start time (45 min post-injection) following the equation: %ID/g = [ROI Uptake (Bq/mm^3^)/Tissue Density (g/mm^3^)] x [1/Injected dose (Bq) x 100].

### Autoradiography

Autoradiography study was performed in C57BL/6 mice (n=2) on day 14 post-K/BxN STA; after the PET imaging, mice were euthanized, and the front and hind limbs were harvested. The tissues were exposed overnight to an E11 phosphor screen and scanned at 50-micron resolution on a GE Typhoon FLA 9500 Variable Mode Laser Scanner. Images of limbs autoradiography were aligned using the “Align Image by Line ROI” function in ImageJ. Fields of low and high ^68^Ga-DOTA-ECL1i signal (2.5 × 2.5 mm) were selected for analysis.

### Assessment of pain-related behaviors

All animals were acclimatized to equipment prior to testing. To assess tactile allodynia, an Electronic Von Frey (EVF) assay was used as previously described [39, 40]. Tactile sensitivity was assessed in both hind limbs using the EVF device (Bioseb) (n=5-8). Pain measurements were conducted prior to K/BxN STA, and at 7-, 14-, and 23-days post-K/BxN STA. The threshold for tactile response was recorded for each measurement. For each limb, three measurements were taken by an individual blinded to treatment, and the results were averaged to provide a representative score for analysis [40].

### Quantitation of proinflammatory mediators

Levels of serum and paw lysate inflammatory mediators including CCL2, IL-6, IL-1β, TNF-α, IFN-γ, and IL-10, were evaluated using BD FACS Calibur Systems (BD Biosciences).

### MicroCT analysis

Following euthanasia, limbs were fixed in 4% paraformaldehyde (PFA) for 24 hours. Fixed limbs were scanned using the Bruker Skyscan 1176 microCT system. Hind and front paws were scanned with a 9-µm isotropic voxel resolution at 50 kV, 500 µA, 980-ms integration time, 3 frame averaging, and 0.5-mm aluminum filter to reduce the effects of beam hardening. Images were reconstructed using NRecon software (with 20% beam hardening correction and 15-ring artifact correction). The outcomes reported in this study include bone mineral density (BMD), bone volume fraction (BV/TV), bone volume (BV), and bone surface-to-volume ratio (BS/BV).

### Histomorphometry and immunofluorescence

Following *ex vivo* microCT scanning, Formalin-fixed paraffin embedded slides of hind paws were prepared at 5 µm thickness, after overnight fixation in 4% paraformaldehyde and decalcification with 10% formic acid (CalEx II). Serial slides were then stained with hematoxylin & eosin (H&E) and toluidine blue (T-blue) according to previous methods [40]. Inflammatory cells infiltrating the synovial lining and the joint cavity were enumerated in 8 to 10 random fields per section using H&E images acquired at 400x magnification. Proteoglycan loss in the cartilage was scored on T-blue–stained sections on a scale from 0 to 3, ranging from fully stained cartilage (score, 0) to fully destained cartilage (score, 3), as previously described [40]. Sections of K/BxN STA hind limbs from CCR2 gfp^+^/^+^ mice (n=4-6/group) on day 7 post-K/BxN STA and serum-negative control tissue were baked (60 °C for 30 minutes) prior to deparaffinization and rehydration with a series of xylene and graded alcohols. Antigen retrieval was implemented with Diva Decloaker 1x in a pressure cooker. To block nonspecific binding, slides received 10% donkey serum in TBS-T (Millipore Sigma) for 1 h at room temperature. Primary antibody (anti-GFP, 1:100 in 1% blocking serum, Cat# NB600-308, RRID:AB_10003058, Novus) was applied overnight at 4 °C, followed by secondary antibody (Donkey antirabbit Cy5, 1:400 in TBS-T, cat# 711-175-152, RRID: AB_2340607, Jackson Immuno Research Laboratories) for 1 h at room temperature the next day. Cell nuclei were labeled with DAPI (1:1000 in PBS for 2 minutes). Slides were washed with tap water in between staining steps and mounted with Diamond ProLong Antifade (ThermoFisher). Digital images of the stained sections were obtained using Leica THUNDER Imager 3D tissue microscope system. Specificity of signals were confirmed using serum-negative controls (processed using the same protocol) and KBN STA negative-controls (processed using the same protocol, without primary antibody application)[42].

### Statistical analysis

Graphing and analysis were performed using GraphPad Prism 10 (GraphPad Software) with a priori significance level (α) set at 0.05. Normality was assessed using Shapiro-Wilk’s test. Mean clinical score for K/BxN mice was determined on the sum of all four limbs. Statistical analyses and the number of animals per group for each experiment are detailed in the respective figure legends. Data are presented as means ± SEM. Comparisons were made using one-way ANOVA, followed by Tukey’s or Dunnett’s post hoc tests, as appropriate.

## Results

### ^68^Ga-DOTA-ECL1i uptake correlates with disease activity and resolution

To define the temporal dynamics of CCR2 expression during inflammatory arthritis, mice underwent serial ^68^Ga-DOTA-ECL1i PET imaging at baseline and at 7, 14, and 23 days following K/BxN STA induction. Quantitative PET analysis demonstrated a rapid increase in tracer uptake that coincided with the onset of clinical arthritis. Uptake within the hind paws increased significantly at day 7 compared with baseline (*P* <0.0001) and remained elevated at day 14 (*P* <0.0001) before declining toward baseline by day 23 (*P* = 0.612) (Figure 1A). In contrast, tracer uptake within the front paws exhibited a transient increase that was significant only at day 7 compared with baseline (*P* <0.0001) and returned to baseline levels thereafter (Figure 1B). These findings indicate that CCR2 expression peaks during the acute inflammatory phase of disease and resolves as arthritis subsides.

Representative maximum-intensity projection (MIP) and fused PET/CT images corroborated the quantitative analyses, demonstrating intense focal accumulation of ^68^Ga-DOTA-ECL1i within the front and hind paws during peak disease, with minimal uptake observed at baseline and following disease resolution (Figure 1C). Increased tracer accumulation was most prominent in the hind paws (PET uptake at day 7 was approximately 2-fold higher than baseline), consistent with the quantitative PET measurements. *Ex vivo* autoradiography further verified localized radiotracer accumulation within inflamed joints, demonstrating high spatial concordance with PET imaging (Figure 1D). Collectively, these findings demonstrate that longitudinal ^68^Ga-DOTA-ECL1i PET sensitively detects dynamic changes in CCR2 expression throughout the progression of inflammatory arthritis.

### Clinical disease severity and pain behavior track longitudinally with ^68^Ga-DOTA-ECL1i uptake

To determine whether longitudinal changes in CCR2 PET reflected disease activity, clinical manifestations of arthritis and pain behavior were assessed throughout disease progression. Ankle thickness increased rapidly following K/BxN STA induction, peaked during maximal disease activity (days 7-9) before steadily declining during disease resolution (Figure 2A). Ankle thickness rose sharply on day 7 (*P* <0.0001), peaked at day 9, remained elevated at day 14 (*P* <0.0001) before returning toward baseline by day 23 (*P* =0.018). Clinical arthritis scores followed a similar temporal pattern (Figure 2B). Following disease induction, clinical scores increased significantly relative to baseline at days 2, 7, 9, and 14 (*P* <0.0001), reaching maximal values during acute arthritis before returning to baseline at day 23 during disease resolution (*P* <0.0001). Likewise, tactile allodynia, measured by the electronic von Frey assay, demonstrated marked mechanical hypersensitivity during active disease (Figure 2C). Withdrawal thresholds were significantly reduced on day 7 compared with baseline (*P* <0.0001) and remained significantly decreased on days 9 and 14 (*P* <0.0001) before partially recovering by day 25 (*P* =0.997).

The temporal concordance between longitudinal PET imaging and clinical outcome measures demonstrates that ^68^Ga-DOTA-ECL1i uptake closely parallels all disease stages, from onset, peak inflammation, and subsequent resolution (Figures 2D-F). PET uptake demonstrated a strong positive correlation with ankle thickness (Pearson *r* = 0.774, *P* < 0.0001) (Figure 2D), confirming that increased tracer accumulation directly reflects degree of joint swelling. Likewise, PET uptake correlated strongly with clinical arthritis scores (Spearman *r* = 0.807, *P* < 0.0001) (Figure 2E), demonstrating that increased CCR2 expression reflects overall disease activity. PET uptake inversely correlated with mechanical hypersensitivity measured by the electronic von Frey assay (Pearson *r* = −0.510, *P* < 0.0001) (Figure 2F), indicating that higher CCR2-mediated inflammation is associated with a lower pain threshold and increased mechanical allodynia.

### Increased ^68^Ga-DOTA-ECL1i uptake in inflamed joints coincided with CCR2^+^ inflammatory monocytes/macrophage accumulation

To define the histopathologic features driving the increased PET signal, we evaluated inflammatory changes within the arthritic joints throughout disease progression. Inflammatory cell influx increased significantly following K/BxN STA induction compared with aged-matched control mice (*P* <0.0001) (Figure 3A), peaking during the acute phase of disease (day 9) and paralleling the temporal increase observed by longitudinal PET imaging. Similarly, cartilage proteoglycan loss tracked with inflammation, peaking at day 9 (Figure 3B).

**Figure 3.**
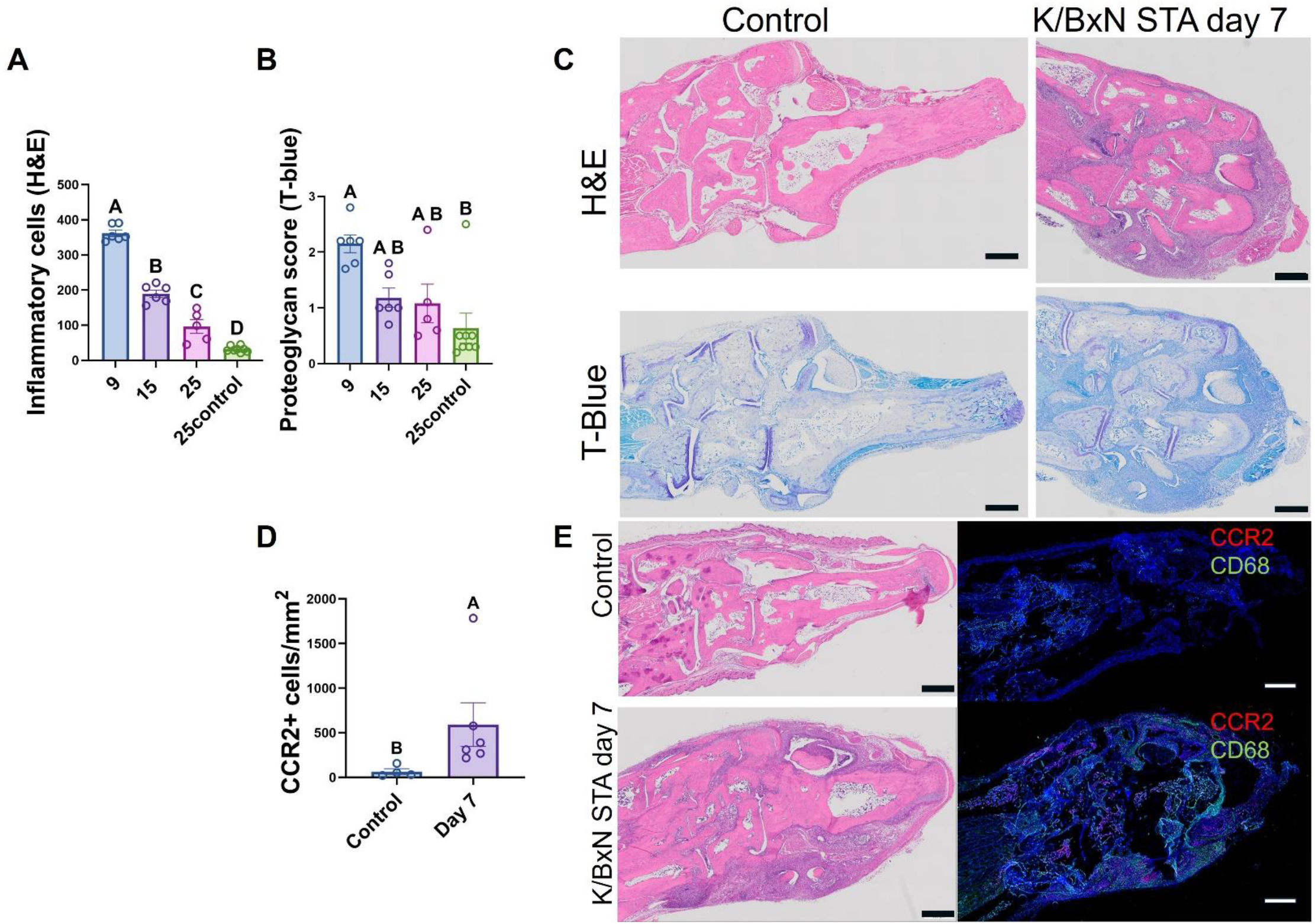
Histopathologic changes and CCR2^+^ cell infiltration during arthritis progression. Histologic inflammation (A) and cartilage proteoglycan loss (B) were quantified in hind paws (*n* = 6–8/group). Representative hematoxylin and eosin (H&E) and Toluidine Blue (T-Blue) sections are shown (C). CCR2^gfp+/+^ mice were used to quantify CCR2^+ cell density (*n* = 5/group), and representative immunofluorescence images of CCR2 (red) and CD68 (green) are shown (D). Data are presented as mean ± SEM. Different letters indicate significant differences (*P* < 0.05) determined by one-way ANOVA with Tukey’s multiple-comparison test (A), Kruskal–Wallis test with Dunn’s multiple-comparison test (B), or unpaired Student’s *t* test (D). Scale bar = 500 μm.

Representative hematoxylin and eosin (H&E) histology revealed marked synovial hyperplasia, inflammatory cell infiltration, and joint tissue expansion in arthritic paws. while Safranin O/Fast Green (Saf-O) staining confirmed progressive loss of cartilage proteoglycan content seven days post-K/BxN STA compared with naïve controls (Figure 3C). These structural changes coincided with the period of highest PET signal, indicating that increased CCR2 expression occurs during active synovitis and early cartilage injury.

To identify the cellular source of the PET signal, we induced K/BxN STA in 6–8-week-old CCR2^gfp+^/^+^ male reporter mice (n=5 K/BxN STA and n=5 naïve controls). Quantitative analysis demonstrated a significant increase in CCR2-positive cells in arthritic joints compared with control mice (*P* <0.0095) (Figure 3D). Immunofluorescence confirmed that CCR2 colocalized extensively with GFP and the resident macrophage marker CD68, identifying infiltrating macrophages as a major driver of increased ^68^Ga-DOTA-ECL1i signal. Together, these findings confirm that elevated PET uptake marks macrophage-rich synovial inflammation driven by infiltration of CCR2-expressing monocytes and macrophages.

To characterize the inflammatory milieu associated with increased CCR2 PET signal, we quantified cytokine and chemokine concentrations in serum and paw lysates throughout disease progression. Distinct temporal patterns were observed between systemic and local inflammatory responses, with the most robust changes observed in the affected joints (Figure 4). Paw lysate cytokine analysis demonstrated a robust local inflammatory response following K/BxN STA induction (Figure 4A–F). CCL2 concentrations increased rapidly, peaking during active disease at day 9 (*P* = 0.0009) before declining during disease resolution (Figure 4A). Similar temporal patterns were observed for IL-1β (*P* = 0.002), IL-6 (*P* = 0.001), IL-10 (*P* = 0.002), and TNF-α (P < 0.0001), all of which reached maximal levels during active disease before decreasing as inflammation resolved (Figure 4B–F). In contrast, IFN-γ levels remained unchanged throughout disease progression (Figure 4E). Furthermore, tracer uptake correlated positively with local CCL2 concentrations measured in paw lysates (Pearson *r* = 0.538, *P* = 0.0004) (Figure 4K), providing direct evidence that the PET signal reflects activation of the CCR2–CCL2 inflammatory axis within arthritic joints.

**Figure 4.**
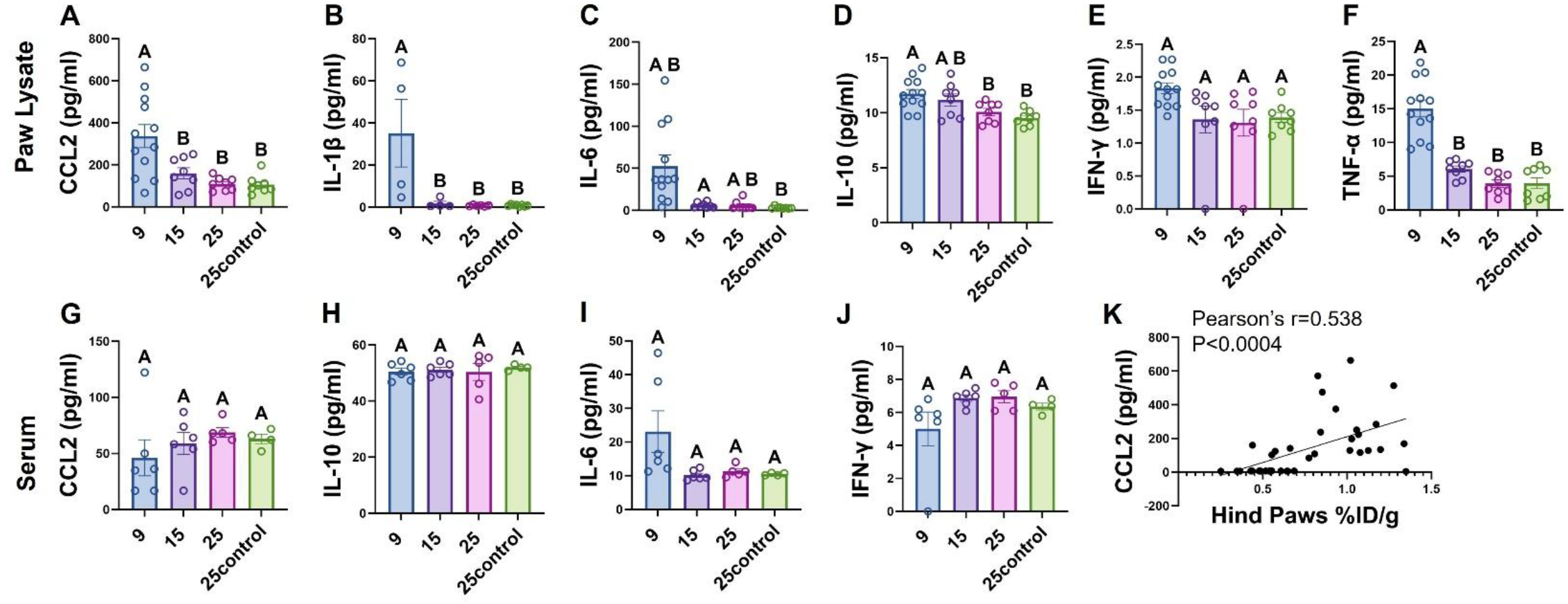
Systemic and local inflammatory mediator profiles during arthritis progression. Serum (*n* = 4– 6/group) and paw lysates (*n* = 4–8/group) were analyzed for CCL2 (A,G), IL-1β (B,H), IL-6 (C,I), IL-10 (D,J), IFN-γ (E,K), and TNF-α (F). Data are presented as mean ± SEM. Different letters indicate significant differences (*P* < 0.05) determined by one-way ANOVA with Tukey’s multiple-comparison test.

Serum cytokine analysis revealed no significant changes throughout disease progression (Figure 4G–J). Serum concentrations of CCL2, IL-10, IL-6, and IFN-γ remained comparable across all time points and control mice, indicating that the systemic cytokine response was not significantly altered despite the pronounced local inflammatory response observed in the paw lysates. The substantially higher cytokine concentrations in paw lysates than in serum indicate that inflammatory activation remained largely localized to the affected joints. Notably, the marked increase in local CCL2 is consistent with the increased recruitment of CCR2-positive inflammatory cells detected by PET imaging and confirmed by immunofluorescence, supporting a localized CCL2–CCR2 inflammatory axis during active arthritis.

### Peak ^68^Ga-DOTA-ECL1i uptake precedes bone remodeling

To determine whether the inflammatory changes identified by ^68^Ga-DOTA-ECL1i PET were accompanied by structural joint damage, microCT was performed. Ex vivo quantitative analysis demonstrated significant alterations in bone architecture in both the hind and front paws compared with control mice (Figure 5). In the hind paws, early bone loss was evident by significant reductions in bone volume fraction (BV/TV) at days 9, 15, and 25 compared with controls (*P* < 0.0001, *P* = 0.005, and *P* < 0.0001, respectively) (Figure 5B) and in bone volume (BV) at days 9 and 25 (*P* = 0.039 and *P* = 0.008, respectively) (Figure 5C). Bone mineral density (BMD) declined later, with a significant reduction on day 25 compared with day 15 (*P* = 0.036) (Figure 5A), indicating progressive deterioration of bone quality over time. Consistent with these findings, the bone surface-to-bone volume ratio (BS/BV) was significantly increased at day 25 (*P* = 0.0006) (Figure 5D), reflecting increased bone surface irregularity and erosive bone remodeling.

**Figure 5.**
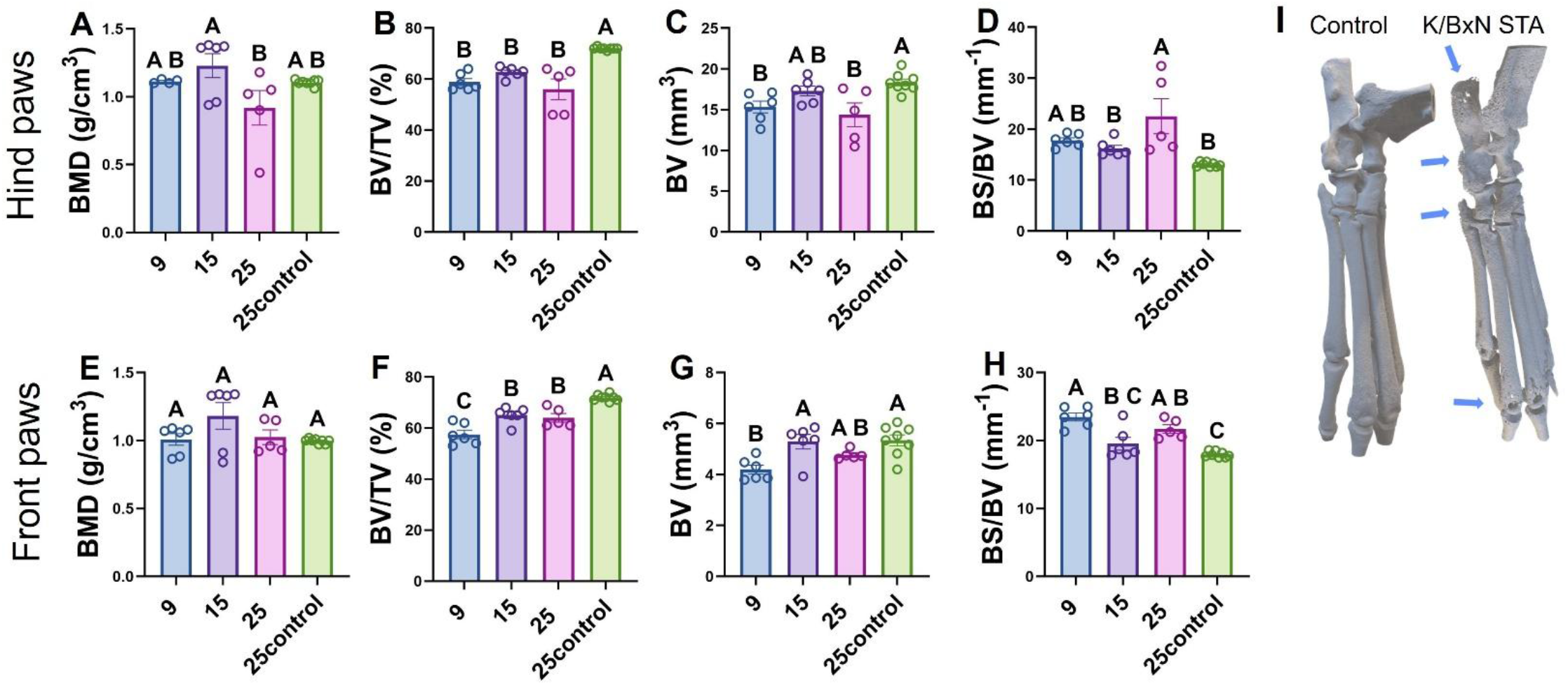
Micro-CT analysis of bone remodeling following K/BxN STA induction. Quantitative microCT analysis of the hind paws (A–D) and front paws (E–H) included bone mineral density (BMD), bone volume/total volume (BV/TV), bone volume (BV), and bone surface/bone volume (BS/BV) (*n* = 4–8/group). Representative three-dimensional reconstructions of control and K/BxN STA hind paws at day 25 are shown (I). Quantitative PET uptake (%ID/g) was correlated with paw lysate CCL2 concentrations (K). Data are presented as mean ± SEM. Different letters indicate significant differences (*P* < 0.05) determined by one-way ANOVA with Tukey’s multiple-comparison test. Pearson correlation analysis was performed.

In the front paws, bone microarchitectural changes were less pronounced than those observed in the hind paws (Figure 5E–H). Bone mineral density (BMD) did not differ significantly among groups (Figure 5E). However, bone volume fraction (BV/TV) was significantly reduced on days 9, 15 and 25 compared with control mice (*P* < 0.0001, *P* = 0.003, and *P* = 0.001, respectively) (Figure 5F). Bone volume (BV) was also significantly decreased on day 9 compared with day 15 (*P* = 0.01) and control mice (*P* = 0.004), whereas day 25 values were not significantly different from either group (Figure 5G). Consistent with these findings, the bone surface-to-bone volume ratio (BS/BV) was significantly increased at day 9 compared with control mice (*P* < 0.0001), indicating early bone surface irregularity and erosive remodeling (Figure 5H).Representative three-dimensional reconstructions further demonstrated cortical irregularities and erosive bone remodeling in arthritic paws at day 25 that were absent in healthy controls (Figure 5I). These findings indicate that the transient increase in CCR2-mediated inflammation detected by PET is followed by measurable structural deterioration of the affected joints.

Collectively, these findings establish ^68^Ga-DOTA-ECL1i PET as a sensitive and quantitative biomarker of CCR2-mediated inflammation in K/BxN STA. Longitudinal PET imaging accurately identified the temporal evolution of inflammatory disease, reflected macrophage-rich synovitis and local CCL2 production, paralleled clinical disease severity and pain behavior, and was associated with subsequent structural joint remodeling. These data support the use of CCR2-targeted molecular imaging as a noninvasive approach for monitoring inflammatory arthritis and quantifying disease activity throughout disease progression.

## Discussion

In this study, we performed a comprehensive longitudinal evaluation of ^68^Ga-DOTA-ECL1i PET imaging to noninvasively visualize and quantify CCR2^+^ inflammatory monocyte and macrophage recruitment during the progression of experimental inflammatory arthritis in the K/BxN serum transfer model. By integrating longitudinal CCR2 PET imaging with histopathology, microCT, and behavioral testing, we establish a comprehensive characterization of the temporal relationship betweenCCR2-mediated inflammation to joint pathology, pain-related behavior, and structural remodeling. Unlike previous imaging studies, which have primarily focused on structural abnormalities, nonspecific metabolic activity, or downstream inflammatory changes, our work directly interrogates CCR2-mediated inflammatory monocyte recruitment, an early pathogenic event in RA. These findings demonstrate that ^68^Ga-DOTA-ECL1i PET serves as a mechanism-based imaging biomarker for early inflammatory arthritis by tracking acute CCR2^+^ inflammatory monocyte and macrophage recruitment. Specifically, we show that CCR2 PET uptake rises sharply during acute arthritis, localizes to macrophage-rich synovium, correlates with clinical disease severity and pain-related behavior, and precedes structural bone damage. Importantly, multimodal validation confirmed that increased tracer uptake reflects localized CCR2^+^ inflammatory cell recruitment within affected joints rather than systemic inflammatory burden. Because CCR2-mediated monocyte recruitment occurs early in RA pathogenesis, this imaging modality could complement existing imaging approaches to identify subclinical inflammation, improve patient stratification, and enhance longitudinal therapeutic monitoring.

A key strength of this study is the comprehensive, multimodal validation of the imaging signal. ^68^Ga-DOTA-ECL1i PET demonstrates strong concordance with autoradiography, histopathology, immunofluorescence, cytokine profiling, pain-related behavior, and microCT. Increased tracer uptake localizes to CCR2^+^ macrophage-rich synovium and correlates strongly with local CCL2 concentrations, supporting activation of the CCL2–CCR2 inflammatory axis. Robust joint inflammation occurs alongside relatively modest systemic cytokine changes, suggesting that CCR2 PET is particularly informative for detecting localized synovial inflammation. Because peak tracer uptake coincides with maximal synovitis and clinical disease activity but precedes structural bone remodeling, our findings support the involvement of CCR2^+^ inflammatory monocyte recruitment in early RA pathogenesis and highlight a potential therapeutic window before irreversible joint damage develops.

Importantly, the correlation between CCR2 PET uptake and pain-related behavior suggests that this imaging approach may provide functional insights beyond inflammatory disease activity, supporting its potential as a molecular imaging biomarker of inflammation-associated pain. This concept is reinforced by a recent clinical study demonstrating an association between increased TSPO PET signal and chronic pain in patients with knee osteoarthritis and chronic low back pain [43]. However, whereas TSPO PET reflects broader immune-related activation, ^68^Ga-DOTA-ECL1i PET selectively targets CCR2-expressing inflammatory monocytes and macrophages, providing greater specificity for a defined proinflammatory cell population implicated in arthritis pathogenesis. Thus, CCR2 PET may offer a complementary and more mechanistically targeted approach to characterize the inflammatory cellular component associated with joint pain. Future studies are warranted to determine whether CCR2 PET can predict pain severity, persistence, or response to anti-inflammatory therapies, supporting its potential translation as a biomarker of inflammation-associated pain.

Beyond established RA, our findings also have important implications for early disease interception. RA develops over several years, during which anti-citrullinated protein antibody (ACPA)-positive individuals may harbor subclinical immune activation before the onset of clinically apparent synovitis. However, serologic markers alone incompletely identify those who will ultimately develop inflammatory arthritis [44], a limitation underscored by recent prevention trials. The StopRA phase II study showed that hydroxychloroquine failed to prevent progression to clinical RA in anti-CCP3-positive individuals [45], while the PRAIRI trial demonstrated that rituximab delayed, but did not prevent, disease onset [46]. Together, these studies highlight the critical need for biomarkers that can detect tissue-level inflammation rather than systemic autoimmunity alone. Given that CCR2-mediated inflammatory monocyte recruitment represents an early pathogenic event, we hypothesize that ACPA-positive individuals may exhibit elevated ^68^Ga-DOTA-ECL1i uptake before clinically apparent synovitis or structural abnormalities become detectable by conventional imaging. If validated prospectively, CCR2 PET could improve patient selection for disease-interception trials, identify individuals at increased risk of developing RA, and provide a quantitative biomarker for monitoring preventive interventions.

Several limitations should be acknowledged. First, the K/BxN serum-transfer model primarily reproduces the innate immune effector phase of inflammatory arthritis and does not fully recapitulate the chronic adaptive immune responses and persistent synovitis characteristic of human RA observed in chronic human disease. Consequently, the transient CCR2 expression observed in this model may differ from that of chronic human RA. Second, only male mice were evaluated. Therefore, future studies should investigate whether sex-specific differences influence CCR2-mediated inflammation and tracer uptake. Although we establish the biological validity of CCR2 PET during disease progression, further investigation is needed to determine whether early changes in tracer uptake predict therapeutic response, long-term structural outcomes, or progression to chronic disease. Future studies should evaluate ^68^Ga-DOTA-ECL1i PET in chronic arthritis models and prospective human cohorts ACPA-positive individuals, patients with clinically suspect arthralgia, treatment-naïve early RA, and established RA to determine whether tracer uptake predicts disease progression and therapeutic response. Integrating CCR2 PET with complementary molecular imaging approaches such as FAPI PET, could further delineate sequential stages of inflammatory monocyte recruitment, fibroblast activation, and tissue remodeling, supporting the development of precision imaging strategies for personalized RA management.

Our findings also address a critical gap in the current imaging landscape in RA. While conventional radiography remains the standard for assessing structural damage, it lacks sensitivity for early inflammation. Ultrasound and MRI substantially improve the detection of early synovitis, bone marrow edema, and erosions [12, 14-18]; however, they primarily characterize tissue-level manifestations of inflammation rather than directly identifying the cellular mechanisms driving disease progression. PET imaging offers an opportunity to bridge this gap. Although ^18^F-FDG PET sensitively reflects metabolic activity, it nonspecifically captures glucose uptake across multiple immune and stromal cell populations [19-21] . Newer targeted PET tracers have improved biological specificity. Translocator Protein Positron Emission Tomography (TSPO) PET reflects broader inflammatory cell activation, including macrophage-associated processes, although its sensitivity for detecting synovitis has been reported to be lower than that of FDG PET and MRI in certain settings [22, 23]. In contrast^68^Ga-FAPI PET primarily maps activated synovial fibroblasts and tissue remodeling [25-28]. Our findings extend these advances by demonstrating that ^68^Ga-DOTA-ECL1i PET captures CCR2-mediated inflammatory monocyte recruitment, an early cellular process implicated in RA pathogenesis. This supports an emerging paradigm in which molecular imaging characterizes distinct cellular mechanisms rather than aggregate inflammatory burden alone.

In conclusion, this study establishes ^68^Ga-DOTA-ECL1i PET as a biologically validated, mechanism-based molecular imaging biomarker capable of noninvasively visualizing CCR2-mediated inflammatory monocyte recruitment during the early stages of experimental inflammatory arthritis. By targeting an early inflammatory cellular process rather than downstream structural consequences, CCR2 PET complements existing modalities while providing mechanistic insights that are not readily obtainable through conventional imaging. Collectively, these findings support the clinical translation of CCR2 PET as a molecular imaging tool with potential applications in detecting subclinical inflammation, identifying individuals at risk of developing RA, monitoring therapeutic responses, and characterizing inflammation-associated pain, ultimately advancing early intervention and precision medicine in rheumatoid arthritis.

## Acknowledgments

The authors would like to thank Nikki Fettig and Margaret Morris for their assistance with the PET/CT scans. Crystal Idleburg and Samantha Coleman from the WashU Musculoskeletal Research Center Histology Core for their assistance in processing the samples. This project was supported by the Mallinckrodt Institute of Radiology at the Washington University School of Medicine; and the Washington University Institute of Clinical and Translational Sciences which is, in part, supported by the NIH/NCATS, CTSA grant #UL1TR002345.

